# The distribution and impact of common copy-number variation in the genome of the domesticated apple, *Malus* x *domestica* Borkh

**DOI:** 10.1101/021857

**Authors:** James Boocock, David Chagné, Tony R Merriman, Michael A Black

## Abstract

**Background:** Copy number variation (CNV) is a common feature of eukaryotic genomes, and a growing body of evidence suggests that genes affected by CNV are enriched in processes that are associated with environmental responses. Here we use next generation sequence (NGS) data to detect copy-number variable regions (CNVRs) within the *Malus* x *domestica* genome, as well as to examine their distribution and impact.

**Methods:** CNVRs were detected using NGS data derived from 30 accessions of *M*. x *domestica* analyzed using the read-depth method, as implemented in the CNVrd2 software. To improve the reliability of our results, we developed a quality control and analysis procedure that involved checking for organelle DNA, not repeat masking, and the determination of CNVR identity using a permutation testing procedure.

**Results:** Overall, we identified 876 CNVRs, which spanned 3.5% of the apple genome. To verify that detected CNVRs were not artifacts, we analyzed the B-allele-frequencies (BAF) within a single nucleotide polymorphism (SNP) array dataset derived from a screening of 185 individual apple accessions and found the CNVRs were enriched for SNPs having aberrant BAFs (P < 1e-13, Fisher’s Exact test). Putative CNVRs overlapped 845 gene models and were enriched for resistance (R) gene models (P < 1e-22, Fisher’s exact test). Of note was a cluster of resistance gene models on chromosome 2 near a region containing multiple major gene loci conferring resistance to apple scab.

**Conclusion:** We present the first analysis and catalogue of CNVRs in the *M*. x *domestic*a genome. The enrichment of the CNVRs with R gene models and their overlap with gene loci of agricultural significance draw attention to a form of unexplored genetic variation in apple. This research will underpin further investigation of the role that CNV plays within the apple genome.

## Background

The availability of genome sequence data from individuals within a species now enables the investigation of a range of inherited genetic variations at a high resolution. Traditionally, genomic analysis of DNA variants has focused on the identification of single nucleotide polymorphisms (SNPs), and small insertions and deletions (INDELS). However in recent years, other forms of genomic variation have also begun to receive attention. One such form is copy number variation (CNV), defined as a deletion, duplication or insertion of DNA sequence fragments longer than 50 base pairs in length [1].

Studies of CNV in eukaryotic organisms, such as dog [2], barley [3], and human [4], have revealed that 4% to 15% of a eukaryotic genome is comprised of regions which exhibit variation in copy number between individuals. CNVs have the potential to exert influence on genes by altering both their expression and structure. For example, in humans the common Sotos syndrome generally occurs when a deletion of one copy of the *plasma coagulation 12* (*FXII*) gene exposes a deficiency in the remaining copy [5]. CNV can also contribute to the phenotypic diversity of domesticated animals: in cattle, a partial deletion of the *ED1* gene causes anhidrotic ectodermal dysplasia [6]; and in pigs, white coat color is caused by a duplication involving the *KIT* gene [7, 8]. CNV distribution is not random across genomes. As an example, segmental duplications (SDs), which are sections of DNA with near-identical sequence, are considered hotspots for CNV formation [9].

CNV is a common feature of plant genomes, and recent work has highlighted its functional relevance. In barley, genes affected by CNV were enriched for potential functions in disease resistance [3], a finding that is consistent with the results of CNV studies in soybean [10] and maize [11]. In wheat, CNVs in *Vrn-A1* and *Ppd-B1* have been shown to influence flowering time [12], and a CNV at the *Rht-D1b* locus is associated with dwarfism [13]. In soybean, increased copy-number of an allele of the *Rhg1* gene is responsible for nematode resistance [14], while in barley, boron tolerance is associated with a CNV at the *Bot1* locus [15]. Based on these reports, there is evidence that CNVs are frequently associated with the biotic and abiotic stress response in annual crop species. However, no analysis of CNVs in perennial tree species has been performed to date. We hypothesize that CNVs are particularly relevant as a source of genetic variability that can be rapidly utilized for adaptation to stress in long-lived plants, including horticultural and forest tree species.

The domesticated apple (*Malus* x *domestica* Borkh.) first appeared in the Near East around 4,000 years ago. In 2010, a high-quality draft genome of ‘Golden Delicious’ was released [16]. Pairwise comparison of the assembled genome revealed that a whole genome duplication (WGD) had recently <50 million years ago occurred in the Pyrinae (which includes fruit species such as apple, pear and quince), leading to an almost doubling of chromosomal number. However, non-global duplication and deletion events, which are localized to individual loci or regions and can thus be considered to represent CNV, have not previously been described in the apple genome. Apple researchers have used Next-Generation Sequencing (NGS) technologies to detect SNPs across the genome, enabling the development of apple SNP arrays used for genomic selection in apple breeding programs and for fine trait mapping [17-19]. NGS-based re-sequencing in the apple genome has indicated that domesticated apple is significantly heterozygous, with 4.8 SNPs per kb, and a recent study has revealed that multiple introgression events with wild apple species such as *Malus sylvestris* and *M. sieversii* have shaped the genomes of modern domesticated cultivars [20].

Hybridization-based methods, such as array comparative genomic hybridization (aCGH), were the first high-throughput approaches used to identify CNVs at the genome-wide scale [21]. More recently however, NGS technologies combined with new analytical approaches, such as read-depth, paired-end, and split-reads analysis, have become popular [22]. Analysis of read-depth is an effective method for CNV detection, relying on detection of changes in the depth of coverage across the genome as being indicative of changes in the underlying copy-number [23]. The aim of our study was to detect CNV regions (CNVRs), in the apple genome using low-coverage (1x to 5x) NGS re-sequencing data from 30 apple varieties grown or used for cultivar breeding worldwide.

## Methods and Materials

### Plant material and Next-Generation Sequencing

A low-coverage NGS dataset for 30 domesticated apple (*M.* x *domestica*) accessions was developed using Illumina GAII technology, with one lane per individual as described in [19]. These varieties represent founders, intermediate ancestors, or important breeding parents used extensively in apple breeding programs worldwide. The complete set maximized coverage of the genetic background of the world’s domesticated apple. Individuals in the set were *Malus* x *domestica* ‘Braeburn’, ‘CrimsonCrisp’, ‘Delicious’, (both the original form and the derived mutant ‘Red Delicious’), ‘Duchess of Oldenburg’, F2268292-2, ‘Geneva’, ‘Golden Delicious’, ‘Granny Smith’, ‘Haralson’, ‘Honeycrisp’, ‘Idared’, ‘James Grieve’, ‘Jonathan’, ‘McIntosh’, ‘Ralls Janet’, ‘Red Dougherty’, ‘Rome Beauty’, ‘Splendour’, ‘Big Red’, ‘Papa Noel’, ‘Merton Russet’, ‘Wilmont Russet’, ‘David’ o.p., ‘Malling 9’, and four advanced selections.

### Sequence alignment and CNVR identification

The reads from the 30 *Malus* x *domestica* samples were aligned to the pseudohaplotype assembly of ‘Golden Delicious’ [16] using the Burrow’s wheeler aligner (BWA v 0.7.5a) maximal exact matches (MEM) command [24, 25]. To reduce the impact of highly abundant organelle DNA on the read depth analysis, a LAST (commit 529:dc2c88c11662) [26] database was created using apple mitochondrial DNA [27] and apple chloroplast DNA [28]. Sections of the assembly with a highly confident match (score >= 40) were considered to be derived from organelles. Additionally, the locations of repeat sequences, such as retrotransposons, were obtained [29] and reads that overlapped repeat sequences, mitochondrial, or chloroplast locations by at least one base pair were removed using bedtools (v2.19.1) [30]. As duplications can occur during the PCR amplification process and create spurious spikes in read-depth signal, these were identified and removed using Picard’s (v 1.124) MarkDuplicates functionality [31].

The binary alignment format (BAM) files used for CNV analysis were filtered based on mapping quality (reads with MAPQ < 20 were discarded), and processed using CNVrd2 [32] (v1.4.0), where the read-depth was counted in 3000-bp non-overlapping windows. Following this, windows that covered a segment of the assembly with greater than 50% Ns were removed. The GC-content of a genomic region influences the read-depth when using Illumina sequencing [33]. This presents a challenge for read-depth analysis and was minimized using the GC-content adjustment method that is implemented in CNVrd2. The resulting GC-adjusted read count data were then standardized within and between samples using CNVrd2. As a result of standardizing across multiple samples, the influence of mappability bias, which is present at some complex regions, was minimized. Coverage statistics were generated using the R programming language (v3.1.1) [34] with figures created in R using the ggplot2 (v1.0.0) [35] and gplots (v2.16) [36] packages. CNVrd2 uses the DNACopy package [37] from the Bioconductor project [38], which employs a binary segmentation algorithm to segment the normalized read-counts. This procedure results in a segmentation score for each window in each sample. These scores represent significant changes in read-depth, with numbers greater than zero indicating a change to increased read-depth, and numbers less than zero indicating a change to decreased read-depth. These changes in read-depth are indicative of changes in the underlying copy-number within a sample. Although these raw segmentation scores can be processed to obtain integer copy-numbers, for this work we chose to analyze the raw segmentation scores, focusing on detecting regions that varied significantly in segmentation score between the apple accessions. Spearman’s correlation was used to investigate the relationship between read-depth and number of segmented regions. A visualization of the distribution of CNVRs throughout the apple was created using Circos (v0.67-4) [39].

To identify CNVRs, a trimmed standard deviation (removing the min. and max. values) was calculated to compare the segmentation scores in each sample for each window. This trimmed standard deviation was used to remove outliers from the calculations. Permutation testing (10,000 permutations per chromosome) was used to determine CNVR significance. This involved shuffling the segmented regions in each sample, recalculating trimmed standard deviations, and calculating an empirical false discovery rate (FDR) [40]. The FDR was calculated at different thresholds, with the genome-wide FDR calculated as a mean of each chromosomal value weighted by chromosome length. A genome-wide trimmed standard deviation threshold of 0.25 was used to determine whether a window was within a CNVR, and neighboring windows above this threshold were merged to form the CNVRs.

### SNP Array Validation of CNVRs

A SNP dataset was used for validation of CNVRs which contained information on the scoring of 10,685 SNP polymorphic markers from the apple Illumina Infinium 20k SNP array [18] screened over 185 apple accessions from the Plant & Food Research germplasm collection that have a similar genetic background to the varieties used for read-depth analysis. SNP genotyping assumes diploid copy number, an assumption that is violated when a SNP falls within CNVRs, as genotypes such as AAB, carrying an extra copy of the “A” allele, and AØ, missing a copy of the “B” allele, may be present [41]. The B allele frequency (BAF), which is the proportion of signal explained by the B allele, has an expected value for heterozygotes of 0.5, and for homozygotes of 0 or 1. Odd numbers of alleles at a locus, such as AAB, may give values falling outside these values. The B allele frequency (BAF) was extracted for each SNP data point using the Illumina GenomeStudio software. CNVRs were tested for enrichment of SNP markers with a B allele frequency (BAF) between 0.05 and 0.35, or 0.65 and 0.95, in at least 10% (19) of samples. Fisher’s exact test [42] was used to check whether the array design was biased against SNPs being located in CNVRs.

### Gene model annotation and functional analysis

The consensus gene models were obtained from the Genome Database for Rosaceae [43]. A gene model was considered to overlap a CNVR if more than 70% of its bases were within the boundaries of a CNVR. Gene Ontology analysis [44] was performed using the Fisher’s exact test. The resulting P-values were adjusted using the FDR-controlling approach of Benjamimi and Hochberg [40]. A separate enrichment test was performed using predicted resistance gene models obtained from the supplementary information of the publication describing the apple genome [16]. The repeat sequences were classified further by BLAST searching [45] against the RepBase19.12 [46] database. The flanking regions of CNVRs were determined by creating a list of positions 10 kb either side of each CNVR, and merging the overlapping regions. The number of elements for each class that entirely overlapped or did not overlap was calculated, followed by enrichment testing using a Fisher’s exact test. Genic regions were determined by creating a list of positions 10 kb either side of each gene model. The average GC content of the flanking regions of both the CNVRs and the genic regions was calculated on a per chromosome basis.

## Results

### Data summary and CNVR identification

Before quality control (QC), the average sequencing coverage for each variety varied between 3.51x and 13.60x. After QC, which included removing reads overlapping the organelle and repeated regions, the average sequencing coverage for each variety was reduced to between 0.95x and 4.96x (Additional file 1), indicating that between 63% and 73% of the reads mapped to the excluded regions.

The read-depth CNV detection method is based on an assumption that the number of reads originating from a region of a genome after removing technical bias is indicative of the copy number for that region. Reads were counted in 3-kb non-overlapping windows. Prior to normalization, the mean number of reads per window varied from 0 to 1911, and the average for all windows was 134 with standard deviation of 103. Following GC-content adjustment and normalization, these windowed read counts were segmented into regions with similar signal values. Investigation of the relationship between coverage and the number of segmented regions revealed a positive correlation (Spearman’s correlation of 0.656) between coverage and the number of regions detected, which is an indication that CNVs are more likely to be detected in samples of higher coverage (Figure 1). Although integer copy number assignment for an individual sample can be performed using the read-depth method, given the low-coverage sequencing data used in our study and the incomplete apple genome assembly, we chose instead to focus on CNVRs that displayed significant variation in segmentation scores and not to attempt integer copy number assignment. These significantly variable CNVRs were determined by summarizing each window using a trimmed standard deviation (removing the minimum and maximum), followed by a permutation testing procedure to calculate the threshold used to identify potential CNVRs: any window with a trimmed standard deviation above this threshold was considered to lie within a CNVR. A threshold of 0.25 gave an acceptable FDR of 11%, and was used in downstream analyses (Figure 2).

**Figure 1.**
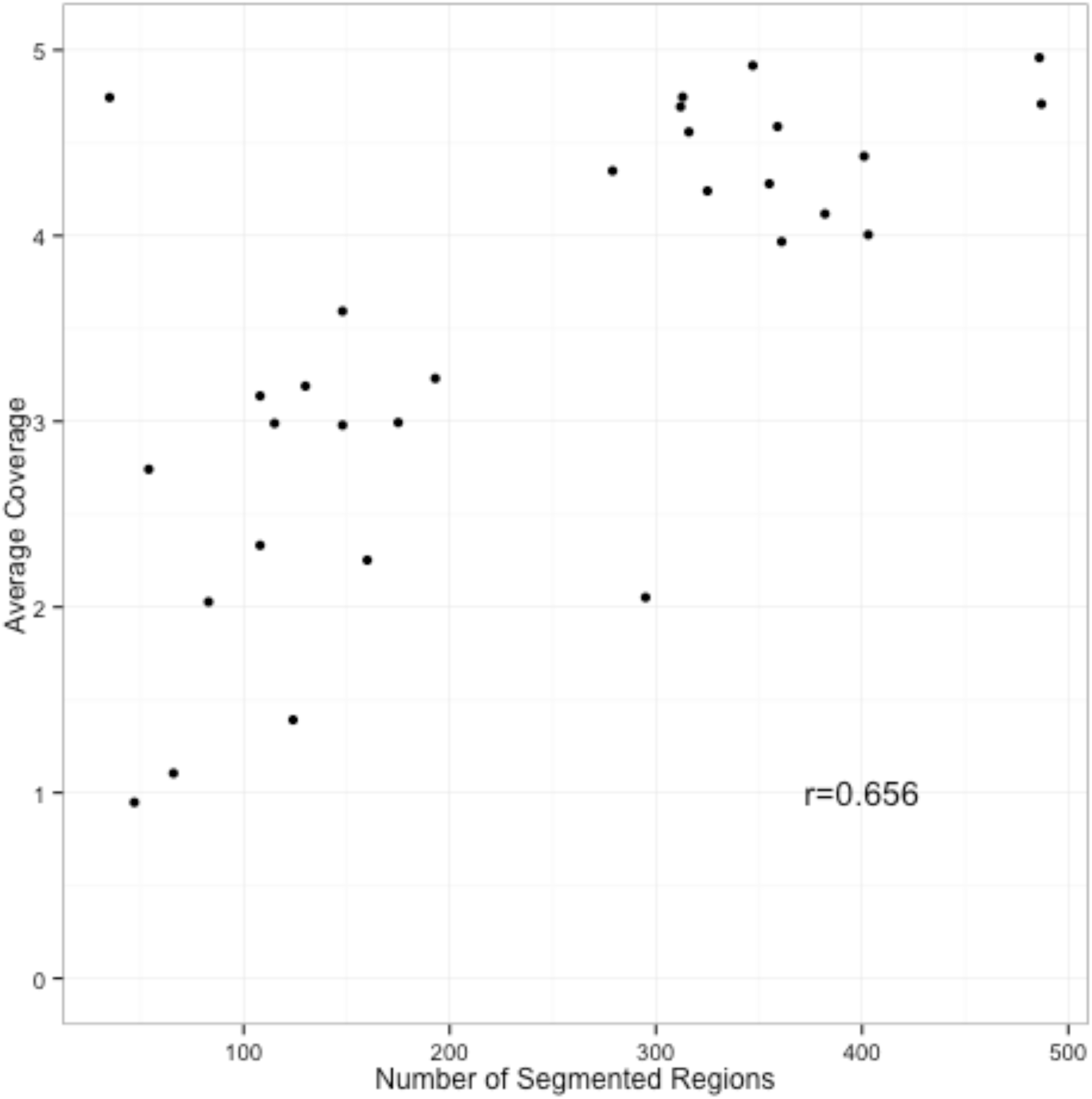
Relationship between number of segmented regions detected in each apple accession and the apple genome coverage. Average coverage per apple accession *versus* total number of segmented regions, determined using CNVrd2.

**Figure 2.**
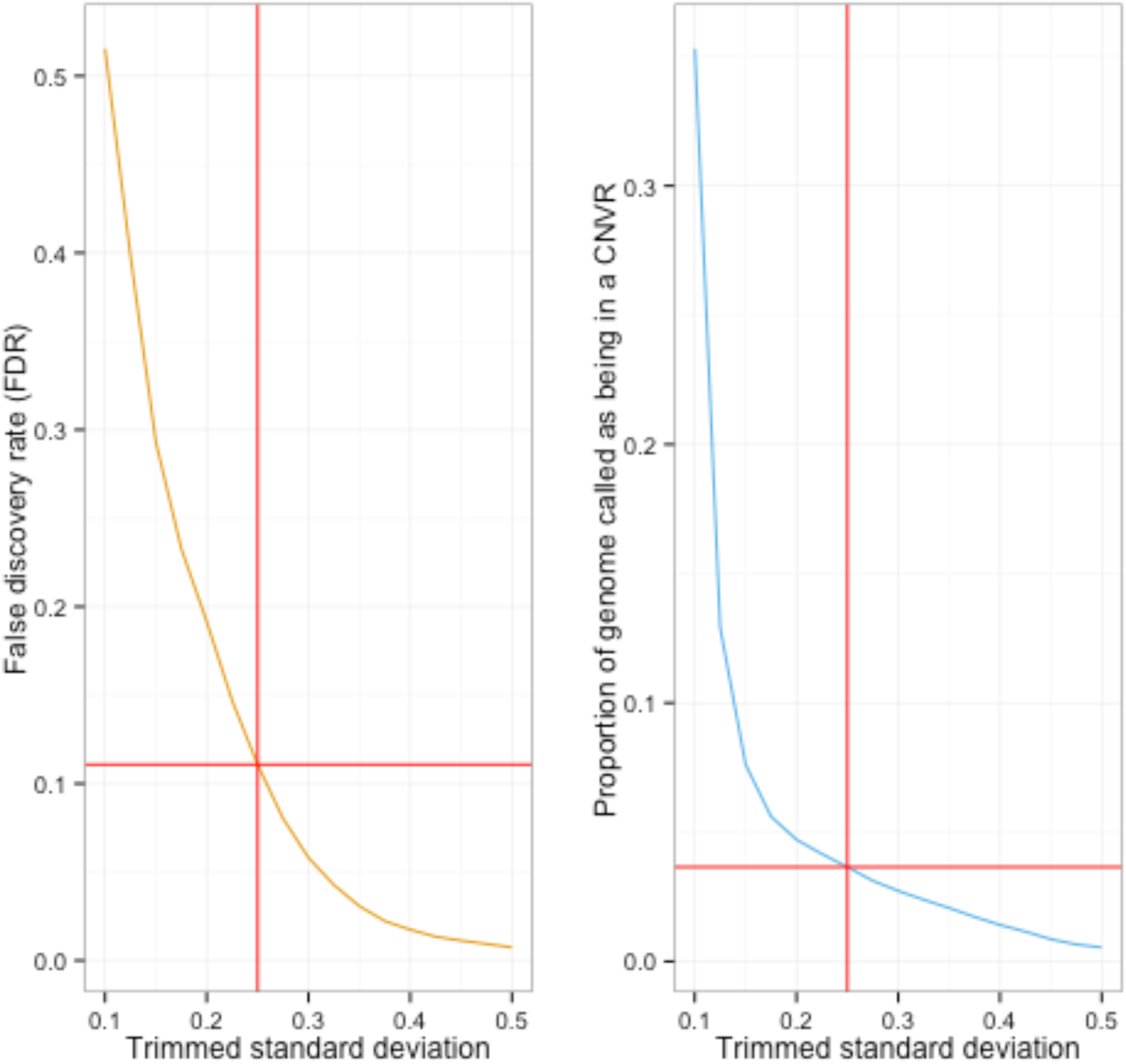
Relationship between the false discovery rate (FDR) estimated using permutation analysis, and the proportion of the genome comprising copy-number variable regions (CNVRs) in the apple genome, as a function of significance threshold. The red lines intersect at a threshold of 0.25, which was the threshold used for downstream CNV analysis using re-sequencing data of 30 apple accessions.

The 876 CNVRs detected using the above threshold spanned a total of 14.4 Mb or 3.5% of the ‘Golden Delicious’ v1.0p pseudohaplotype assembly (Figure 5; Additional file 2). They ranged in size from 3kb to 99kb, with an average length of 16.4kb, and a median length of 12kb (Figure 3). CNVRs appeared to be non-randomly distributed throughout the genome (Figure 4). The percentage of an individual chromosome in CNVRs varied between 1.5% for chromosome 16 and 5.7% for chromosome 10 (Additional file 3). When the removal of regions split a large CNVR, it was considered as two smaller CNVRs.

**Figure 3.**
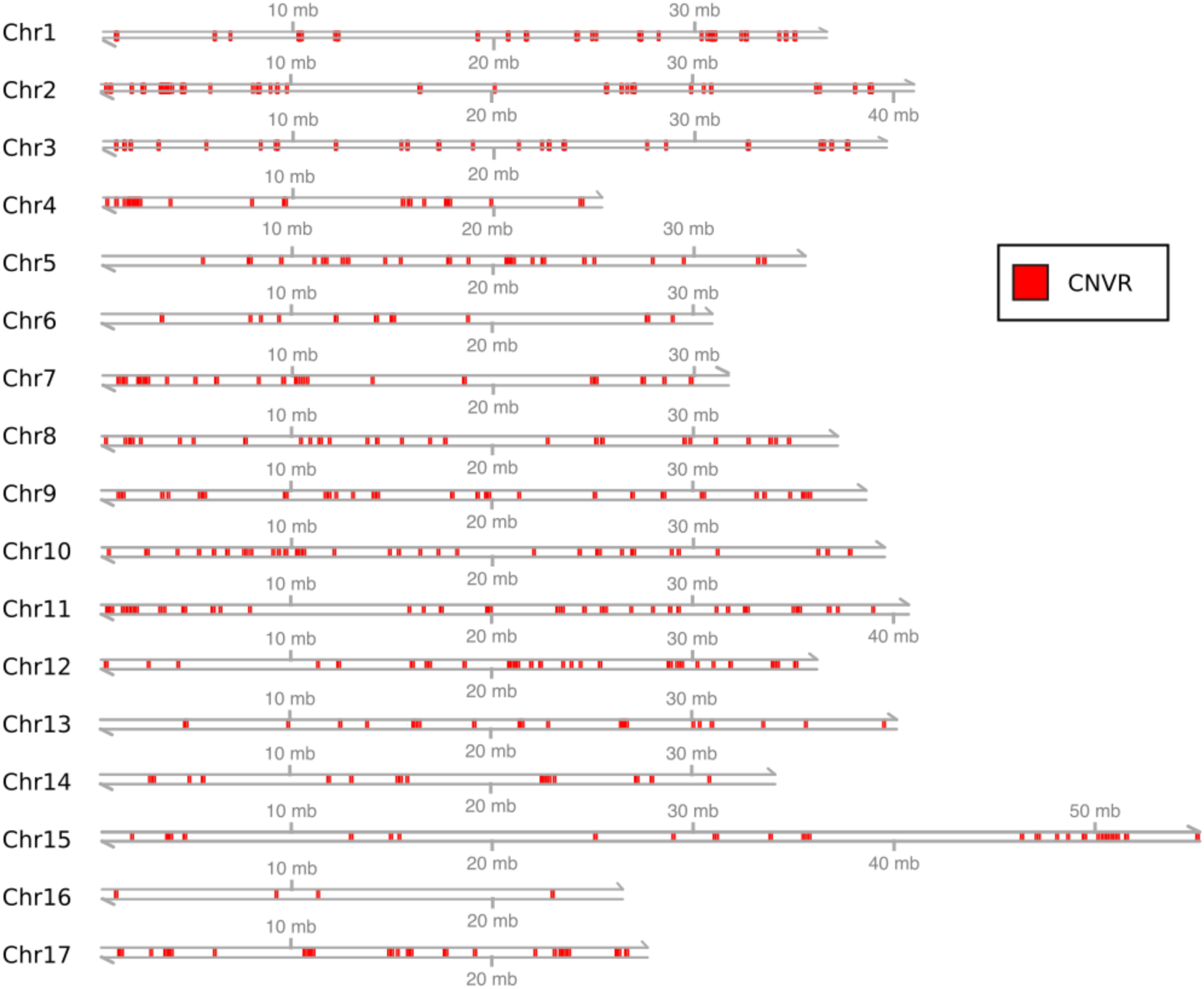
Copy-number variable regions (CNVRs) distribution in the apple genome. The 17 grey lines represent all the chromosomes of the ‘Golden Delicious’ v1.0p pseudohaplotype assembly [16]. Red sections indicate the locations of the 876 CNVRs. mb: megabases.

**Figure 4.**
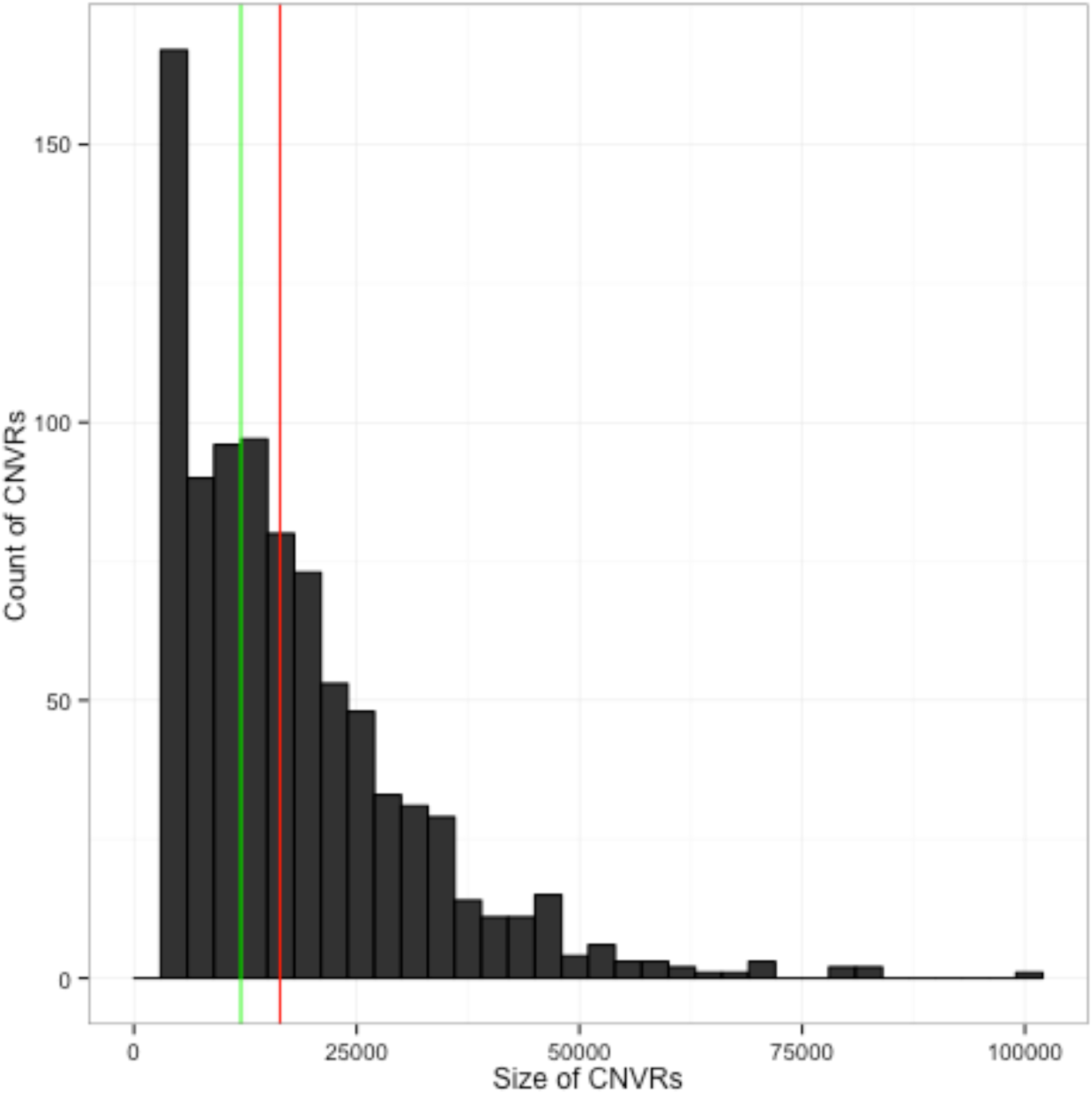
Distribution of length and frequency of copy-number variable regions (CNVRs) in the apple genome. Size (in base pairs) versus number of CNVRs. The green and red lines represent the median and mean CNVR size, respectively.

**Figure 5.**
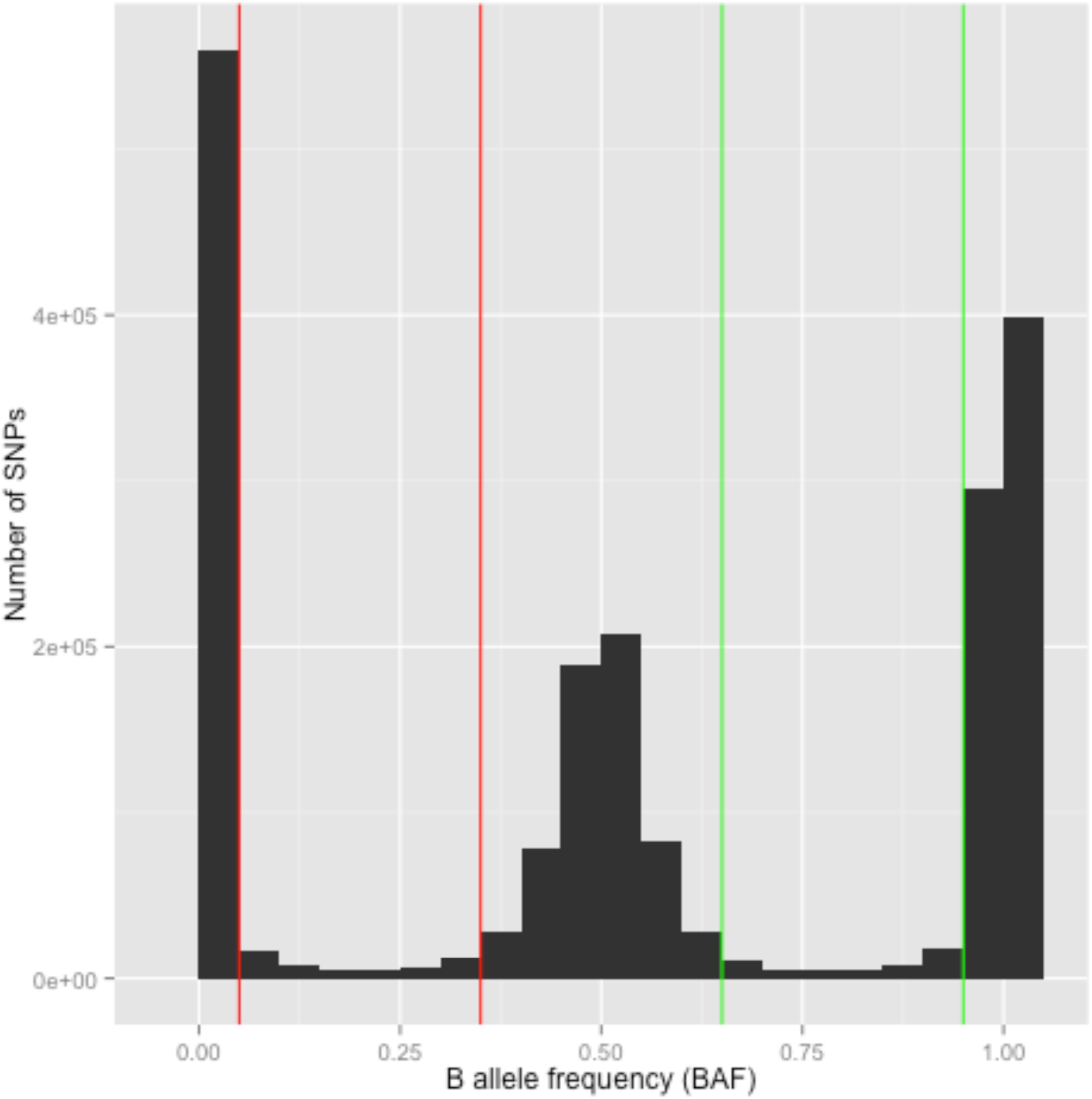
Distribution of B allele frequencies for the 20k SNP array screened on 185 apple accessions. The count of all B allele frequencies (BAFs) is displayed in windows of size 0.05 from a dataset containing 185 apple accessions genotyped on the apple Infinium 20k SNP array [18]. The ranges 0.05-0.35 and 0.65-0.95, shown by red and green lines respectively, indicate the ranges where an individual BAF was considered aberrant.

### Repeat Analysis and GC Content

Repeated sequences were investigated for their relationship with CNVRs. A list of 10-kb regions flanking CNVRs was developed, with overlapping sections merged. The repeated elements that are the most numerous in the apple genome (*Copia*, *Gypsy*, *hAT*, *Cassandra*, and LINE) [16] were tested for enrichment within the flanking regions. A significant depletion was observed for *Gypsy* elements (P = 0.007, Fisher’s exact test) and a significant enrichment for *Copia* elements (P = 0.006, Fisher’s exact test) (Table 1). No significant enrichments or depletions (P > 0.05, Fisher’s exact test) were observed for *hAT*, Cassandra, and LINE elements, and no overall enrichment (P > 0.05, Fisher’s exact test) was detected for repeated elements. The genome-wide GC content of CNVRs was nominally (P = 0.03, T-test) higher (average 37.9%) than that of the pseudohaplotype assembly (37.8%). In contrast, the genome-wide average for genic GC content was nominally (P = 0.02, T-test) lower than that of CNVRs (average 37.6%) (Additional file 3).

**Table 1:** Number of repetitive elements contained in the 10 kb of sequence up- and downstream of the copy-number variable regions (CNVRs) in the apple genome.

| CNVR flanking Regions |  |  | non-CNV flanking regions |  |  |
| --- | --- | --- | --- | --- | --- |
| Repeat Type | Observed | Expected | Observed | Expected | P-value |
| *Gypsy* | 1917 | 2087.5 | 85346 | 85175.5 | 0.007 |
| *Copia* | 815 | 707.7 | 28767 | 28874.3 | 0.006 |
| *Cassandra* | 245 | 225.2 | 9169 | 9188.8 | 0.375 |
| *LINE/RTE* | 264 | 251.5 | 10250 | 10262.5 | 0.624 |
| *hAT* | 402 | 350.6 | 14253 | 14304.4 | 0.065 |
| Total | 3643 | 3622.4 | 147785 | 147805.6 | 0.812 |

### Validation of CNVRs using a SNP array

Although low density SNP microarrays do not enable copy-number detection directly, because of their limited probe density, they do present a method of independently validating CNVRs detected by read-depth analysis. We accomplished this by extracting the BAFs from an apple Illumina Infinium 20-k SNP array dataset that contained the genotyping information for 185 accessions. After the removal of SNPs that overlapped the windows which were previously removed from the analysis because they contained Ns, repeats, or organelle DNA, 10,685 SNPs remained. BAF thresholds of 0.05-0.35 and 0.65-0.95 were considered aberrant (Figure 4) and when 19 (10%) of samples fell within these ranges, the SNPs were considered as having an aberrant BAF pattern indicative of CNV. CNVRs contained 49 of the 1845 (2.6%) SNPs with aberrant BAFs, and 115 of the 8840 (1.3%) normal (disomic) SNPs, representing an approximately two-fold enrichment of SNPs with aberrant BAFs, within CNVRs (P < 0.0001, Fisher’s Exact test). Interestingly, 98.5% of the SNPs represented on the apple Illumina Infinium 20k SNP array were located in non-CNV regions, indicating a significant depletion of SNPs in CNVRs (P < 1e-13, Fisher’s Exact test).

### Functional Analysis of CNVRs

Of the consensus gene models from the Genome Database for Rosaceae, 29,015 were located in genomic regions that were analyzed. A total of 845 (2.9%) of these gene models exhibited a minimum of 70% of their base pairs overlapping putative CNVRs (Additional file 4). Significant depletion of gene models within CNVRs was observed (P < 1e-6, Fisher’s Exact test), with the proportion of the genome assembly spanned by CNVRs (3.5%) being greater than the proportion containing gene models. Over half (470) the CNVRs did not contain a gene model, with the remaining 406 regions (46.3%) containing at least one gene model. Functional enrichment analysis of CNVRs revealed 20 GO terms overrepresented after FDR p-value adjustment (Table 2). These terms included “apoptotic process”, “innate immune response” and “defense response”, for which a significant proportion of annotations were found to originate from resistance (R) gene models. This class of genes are proteins containing nucleotide-binding sites (NBS or NBC-ARC domains) and C-terminal leucine-rich repeats (LRR), and are key components of the immune response in plants [47, 48]. To investigate the relationship between R gene models and CNVRs directly, we obtained a list of 992 resistance (R) gene models from the apple genome publication [16]. Of the 992 R gene models, 268 were located in the GD pseudohaplotype assembly we used for CNV detection, and 47 (16.3%) of these R gene models overlapped CNVRs (Additional file 5). Furthermore, R gene models were enriched greater than five-fold within CNVRs (5.5% of total gene models) compared with outside these regions, where they comprised 0.8% of total gene models (P < 1e-22, Fisher’s exact test) (Table 3). Ten of these enriched GO terms were not attributable to R gene models, such as “ion transport”, “chloride transport”, and “voltage-gated chloride channel activity”.

**Table 2:** Significant Gene Ontology terms (P<0.05 after False-discovery rate correction) for gene models located within copy-number variable regions (CNVRs) in the apple genome.

| **Geno Ontology ID** | **Description** | **P value** |
| --- | --- | --- |
| GO:0006915 | apoptotic process | 4.23E-48 |
| GO:0031224 | intrinsic component of membrane | 4.09E-43 |
| GO:0004888 | transmembrane signaling receptor activity | 1.16E-42 |
| GO:0045087 | innate immune response | 2.75E-42 |
| GO:0005515 | protein binding | 1.03E-32 |
| GO:0007165 | signal transduction | 8.93E-31 |
| GO:0005524 | ATP binding | 9.26E-22 |
| GO:0006952 | defense response | 2.40E-21 |
| GO:0005216 | ion channel activity | 0.006 |
| GO:0006508 | proteolysis | 0.009 |
| GO:0004674 | protein serine/threonine kinase activity | 0.011 |
| GO:0006468 | protein phosphorylation | 0.011 |
| GO:0043086 | negative regulation of catalytic activity | 0.011 |
| GO:0005516 | calmodulin binding | 0.012 |
| GO:0005247 | voltage-gated chloride channel activity | 0.026 |
| GO:0042802 | identical protein binding | 0.028 |
| GO:0004497 | monooxygenase activity | 0.042 |
| GO:0006821 | chloride transport | 0.042 |
| GO:0030552 | cAMP binding | 0.045 |
| GO:0016710 | trans-cinnamate 4-monooxygenase activity | 0.049 |

**Table 3:** Resistance (R) gene models located within copy-number variable regions (CNVRs) in the apple genome.

| MPIDs | Chromosome | Start (bp) | End (bp) | Width (bp) |
| --- | --- | --- | --- | --- |
| MDP0000816743 | chr1 | 24886957 | 24890824 | 3867 |
| MDP0000144874 | chr1 | 30626140 | 30628752 | 2612 |
| MDP0000634037 | chr1 | 30801610 | 30806248 | 4638 |
| MDP0000257201 | chr1 | 30817742 | 30823513 | 5771 |
| MDP0000282740 | chr1 | 30901618 | 30906893 | 5275 |
| MDP0000149577 | chr1 | 31011253 | 31015489 | 4236 |
| MDP0000685087 | chr2 | 2694102 | 2698622 | 4520 |
| MDP0000483490 | chr2 | 2705247 | 2706068 | 821 |
| MDP0000577795 | chr2 | 3531631 | 3532194 | 563 |
| MDP0000478441 | chr2 | 3606656 | 3610584 | 3928 |
| MDP0000522693 | chr2 | 3615408 | 3616082 | 674 |
| MDP0000563856 | chr2 | 3870168 | 3872449 | 2281 |
| MDP0000466397 | chr2 | 3940831 | 3943960 | 3129 |
| MDP0000458576 | chr2 | 3967880 | 3972931 | 5051 |
| MDP0000746482 | chr2 | 3985869 | 3990107 | 4238 |
| MDP0000460947 | chr2 | 4615319 | 4619759 | 4440 |
| MDP0000512349 | chr2 | 4701149 | 4701745 | 596 |
| MDP0000573272 | chr2 | 4731508 | 4735373 | 3865 |
| MDP0000206582 | chr2 | 9335438 | 9338647 | 3209 |
| MDP0000301892 | chr2 | 9357524 | 9362451 | 4927 |
| MDP0000640508 | chr2 | 38880821 | 38885458 | 4637 |
| MDP0000261720 | chr2 | 38910867 | 38912498 | 1631 |
| MDP0000303694 | chr3 | 22748766 | 22753650 | 4884 |
| MDP0000173893 | chr4 | 15497478 | 15501779 | 4301 |
| MDP0000399716 | chr6 | 15009829 | 15020552 | 10723 |
| MDP0000148432 | chr6 | 15038138 | 15040970 | 2832 |
| MDP0000257606 | chr6 | 15042847 | 15050886 | 8039 |
| MDP0000166759 | chr7 | 1337480 | 1340290 | 2810 |
| MDP0000196621 | chr7 | 1370821 | 1373441 | 2620 |
| MDP0000259862 | chr7 | 1591348 | 1594205 | 2857 |
| MDP0000493661 | chr7 | 2442347 | 2446032 | 3685 |
| MDP0000321189 | chr7 | 2458873 | 2466023 | 7150 |
| MDP0000685425 | chr8 | 11405302 | 11410962 | 5660 |
| MDP0000586712 | chr9 | 1526915 | 1531337 | 4422 |
| MDP0000125338 | chr9 | 19770884 | 19773602 | 2718 |
| MDP0000472864 | chr10 | 892484 | 896484 | 4000 |
| MDP0000264448 | chr10 | 10432552 | 10452135 | 19583 |
| MDP0000193087 | chr11 | 1928588 | 1932682 | 4094 |
| MDP0000732188 | chr11 | 2110384 | 2116572 | 6188 |
| MDP0000212932 | chr11 | 3506106 | 3509841 | 3735 |
| MDP0000230932 | chr11 | 3712888 | 3720930 | 8042 |
| MDP0000171723 | chr13 | 21556571 | 21559228 | 2657 |
| MDP0000168665 | chr14 | 13071029 | 13075189 | 4160 |
| MDP0000272830 | chr15 | 15393697 | 15404409 | 10712 |
| MDP0000680657 | chr15 | 48113339 | 48116942 | 3603 |
| MDP0000312626 | chr17 | 1585061 | 1588664 | 3603 |
| MDP0000291677 | chr17 | 17704304 | 17710114 | 5810 |

### CNVR co-location with apple QTLs

A number of quantitative trait loci (QTLs) linked to a range of fruit, pest and disease resistance and tree physiology characters have been mapped in apple. The location of CNVRs was compared to the location of mapped QTLs, with a focus on adaptive traits such as response to biotic stress, phenology and fruit quality.

A QTL region linked to bud phenology has previously been mapped at the top of chromosome 9, with some positional candidate genes proposed. Three CNVRs overlap with this QTL region at positions 1,404,000 to 1,671,000 bp on chromosome 9. A candidate gene (MDP0000241327) that encodes for a putative *E3 ubiquitin-protein ligase*, which is duplicated 14 times in tandem and located within the bud phenology QTL region, is surrounded but not included in these CNVRs [49].

*Pl2* is a major resistance gene to powdery mildew (*Posdosphaera leucotricha*), derived from *Malus zumi* [50]. A CNVR on chromosome 11 contains the likely location of the marker FBsnPl2-2, which co-segregates with this trait. This CNVR is located at positions 3,506,106 to 3,509,841 and contains a predicted R gene (MDP000021232) [51].

Resistance loci to apple scab (*Venturia inaequalis*) derived from a range of apple cultivars and wild germplasm accessions have been mapped to chromosome 2. Nineteen CNVRs are found within an interval between positions 3,507,000 bp and 4,107,000 bp on the primary assembly of ‘Golden Delicious’ v1.0p. Seven R genes (MDP000057795, MDP0000478441, MDP0000522693, MDP0000563856, MDP0000466397, MDP0000458576, and MDP0000746482) are found within this cluster of CNVRs (Figure 6 & Figure 7). Sequences linked to *Rvi4/Vh4* [52] (KM105054) and *Rvi15/Vr2* [53] (KM105120) for which genetic markers tightly linked to these resistances have been developed are located within this region [51].

**Figure 6.**
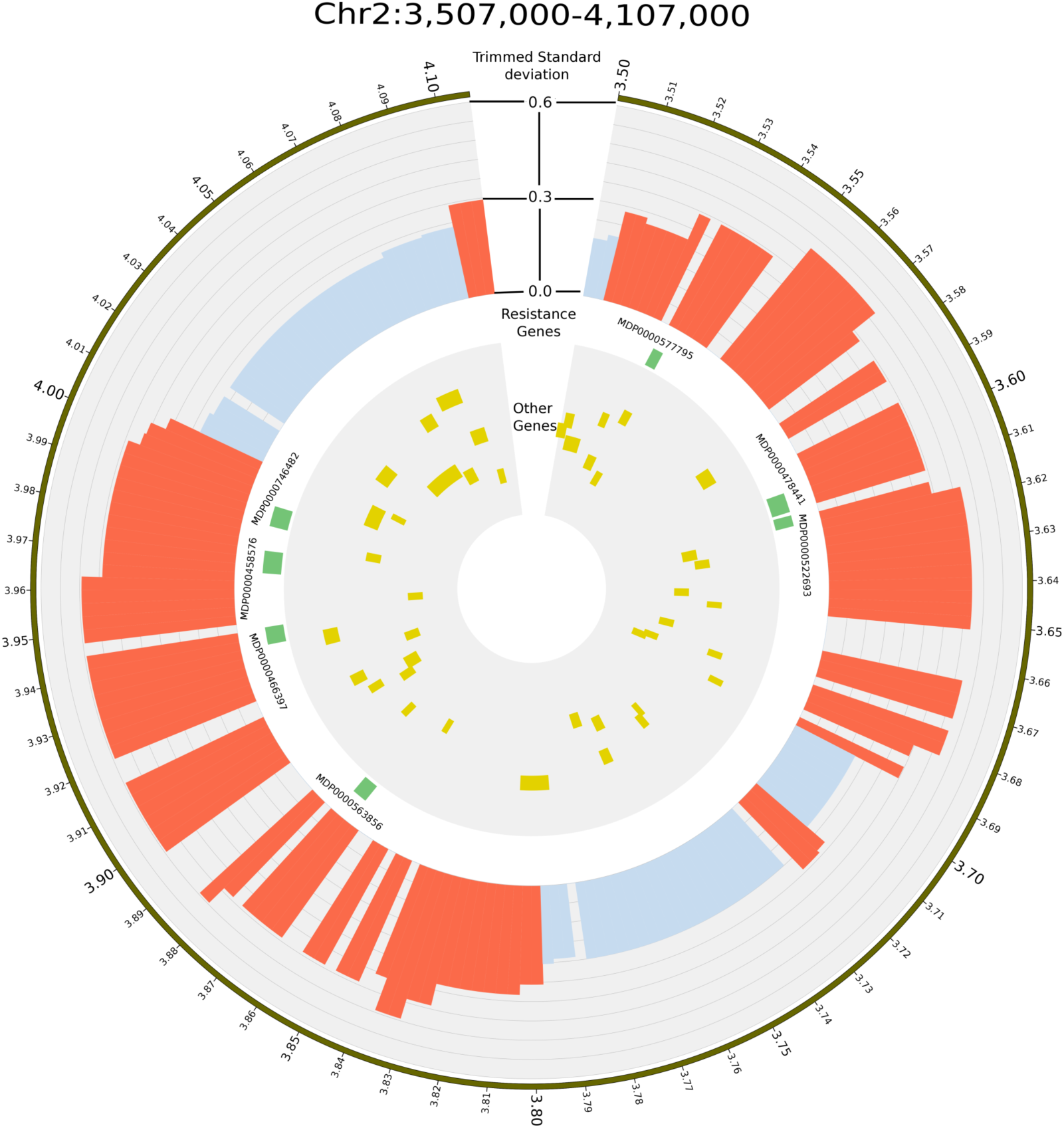
Circos plot displaying the copy-number variable regions (CNVRs), R genes, and genes in the region between positions 3,507,000 bp and 4,107,000 bp on apple chromosome 2. The figure shows the region between 3,507,000 bp and 4,107,000 bp on chromosome 2 based on the ‘Golden Delicious’ v1.0p pseudohaplotype assembly [16]. This region contains resistance loci to apple scab, including *Rvi4/Vh4* and. *Rvi15/Vr2.* Tracks from outside to inside are: (1) region ideogram ticks show the position in megabases (Mb), (2) trimmed standard deviation of segmentation scores plotted as a histogram with a y-scale from 0 to 0.6 (red bars indicate regions designated as CNVRs, blue bars represent all other regions), (3) the location of resistance genes, and (4) the location of genes other than resistance genes (yellow bars represent the location of these genes).

**Figure 7.**
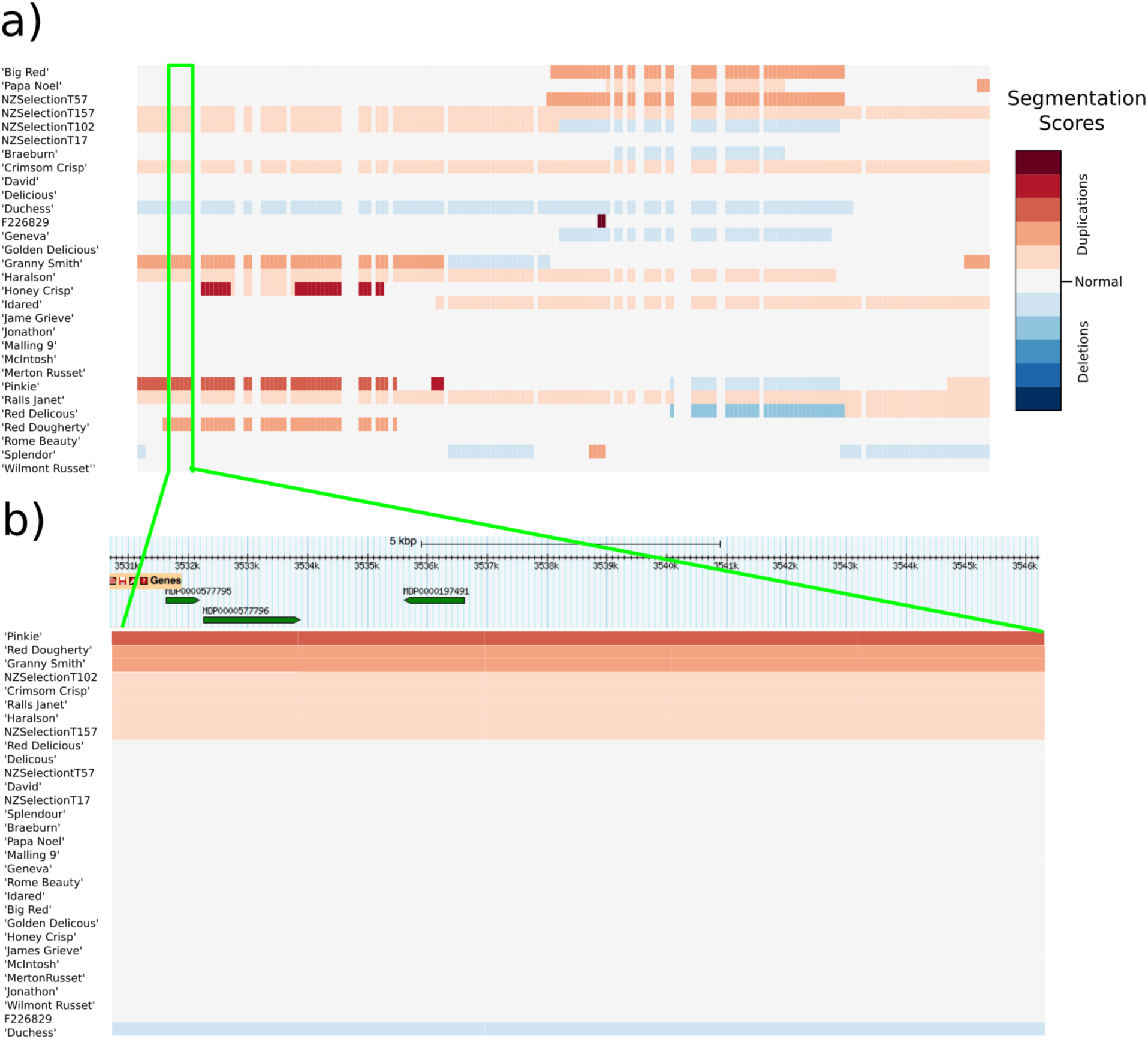
Heatmap representation of the segmentation scores of the region between positions 3,507,000 bp and 4,107,000 bp on apple chromosome 2. (a) Heatmap of segmentation scores with rows ordered in alphabetical order of accession name for the region chromosome 2 from 3,507,000 bp to 4,107,000 bp. The segmentation values are plotted for each window in the region; these values represent significant changes in read-depth signal. The green highlighted region shows one of the 19 CNVRs located within the region. (b) Heatmap of segmentation scores with rows ordered by segmentation score for the CNVR on chromosome 2 between positions 3,531,000 bp and 3,546,000 bp. The resistance gene model (MDP000057795) and the other gene models (MDP000057796 and MDP0000187491) are located above the heat map in their correct position relative to their location within the CNVR. The track above the heatmap displays the location of the gene models found within the CNV (extracted from the Rosaceae genome browser).

## Discussion

Characterization of the apple genome has focused primarily to date on whole genome duplication, the detection of SNPs, the prediction of genes and the characterization of gene families, as well as generation of an inventory of repeated elements. Structural variations, such as copy-number, have not been studied in detail, in apple or indeed in other fruit tree species. In short-lived crop species, such as maize and rice, copy-number variation has been investigated using NGS and microarray technologies, and a growing consensus has emerged as to the agricultural relevance of CNV [54]. In particular, genes located in CNVRs have been observed to be enriched for stress-related response genes, including the canonical R-genes, which contain a leucine rich-repeat (LRR) domain. This highlights the importance of CNV in relation to agricultural crop genetics, and the need to characterize this type of variation in horticultural species. In this study, we used NGS data from 30 apple accessions to identify 876 copy-number variable regions (CNVRs). The CNVRs within the apple genome overlap 845 gene models and are enriched for R gene models. A significant enrichment of SNPs with aberrant BAFs was observed within CNVRs by using a SNP derived from a screening of 185 individual apple accessions with similar genetic background to those of the 30 varieties used for read-depth analysis. This strengthens the evidence that the regions we detected represent true CNVRs.

Although CNV had not been investigated previously in the apple genome, there has been some evidence that CNV might play an important biological role here. A study investigating QTLs associated with bud phenology in apple identified a candidate *E3 ubiquitin protein* encoding gene present as a tandem array of 14 copies in ‘Golden Delicious’, making it a putative CNV [49]. More recently, a study that was performed to design SNP markers for eight major disease resistance loci, encountered two problems that can be explained by the occurrence of CNV in these regions [51]. Firstly, for the *Rvi2*, *Rvi4* and *Rvi11* loci conferring resistance to apple scab (*Venturia inaequalis*), the presence of paralogs made it difficult to design primers for the SNPs linked to the respective loci, which amplified alleles only at the specific region of interest, without co-amplifying the paralogous sequences. Paralogs located in tandem would be considered a segmental duplication, which are hotspots for CNVs [9]. Secondly, SNPs linked to *Pl2*, a major locus conferring resistance to powdery mildew (*Posdosphaera leucotricha*), failed to amplify when resistance was not observed. This would be expected if resistance were conferred by the insertion of a segment of DNA, which contains the SNPs that failed to amplify, and is absent from susceptible individuals [51].

### Data QC and analysis approach

Because of apple’s high heterozygosity, its genome was not assembled as a single reference sequence but rather into four haplotypes, or individual sequences. The primary pseudohaplotype assembly (*Malus* x *domestica* Whole Genome v1.0p) [55], which was used in this analysis, represents the shortest path through the sequence scaffolds. This reference was used to facilitate short-read mapping and read-depth analysis, using a workflow analogous to studies in diploid mammals, for which many bioinformatic tools have been developed [22]. Repeated elements, which are estimated to cover 42.4% of the apple genome [16], create a problem for read-depth analysis, as reads which map to repeat regions of the reference create noise in the read-depth signal. This interferes with data normalization and subsequently the downstream segmentation procedure. Many CNV studies attempt to correct for this effect by masking repeat regions before alignment (i.e., changing the position in the reference to an ‘N’) [56, 57]. However, as the quality score of reads is determined by the uniqueness of the mapping position, masking may lead to reads that originate from the repeated sequences mapping spuriously to other locations, with potentially high quality scores. In our analysis, reads were first mapped, and only then were repeated elements removed. This represents an alternative and potentially improved approach to overcoming the problem of repeated sequences. The presence of chloroplast and mitochondrial DNA within the reference genome creates a similar issue, as the large number of these organelles in each cell would lead to large numbers of reads mapping to these regions, potentially interfering with the read-depth approach. In the analysis presented here, reads mapping to organelle-derived regions were removed before CNV segmentation.

Low-coverage NGS re-sequencing data were used to identify the CNVRs that exhibited significant variation in copy-number within a set of apple accessions with genetic links to international breeding programs. The CNVRs identified were found to vary greatly in copy-number among the accessions, and represent candidate gene loci that may be involved in control of variability in traits, such as pest and disease resistance. The number of samples used (30 apple accessions) is higher than employed in some recent read-depth studies in domesticated organisms [e.g., the number of samples was five and 16 in CNV studies of cows [57] and pigs, respectively [56]]; however, is smaller than studies in maize [58] and soybean [10], which involved 278 and 302 samples, respectively. The average coverage for sequencing of our samples ranged from 0.95x to 4.96x, which is relatively low and might have introduced noise into the read-depth signal. Our finding that a strong correlation was observed in our dataset between sequencing coverage and the number of detected CNVs suggested that the read-depth method is more efficient at detecting copy-number changes in higher-depth samples, and as low-coverage data do not enable reliable calling of exact copy number at specific loci for individuals, we opted to analyze the variation among segmentation scores across the 30 varieties to provide an inventory of CNVRs in the apple genome. A trimmed standard deviation was chosen (removal of the min. and max. values), to reduce the impact of outliers potentially driven by noise, and a permutation test was used to assess the likelihood of finding such a variable region by chance. This allowed us to determine a conservative threshold for CNVR identity, with what we considered an acceptable FDR of 11% [40].

### Patterns of CNVRs in the apple genome

The read-depth analysis identified 876 CNVRs in the apple genome, ranging in length between 3 kb and 99 kb, with an average of 18 kb. It should be noted that because the CNVRs separated by windows that were removed were not merged, the number of detected CNVRs is likely to be an overestimate, as a number of CNVRs interleaved with removed regions may actually comprise a single large CNVR. Our results suggest that 3.5% (14.4Mb) of the analyzed assembly lies within a CNVR. This percentage is lower than found in previous CNV studies in plants; for example, the CNV percentages found in maize [58] and barley [3] were 10% and 14.9%, respectively. This discrepancy might be explained by the lower degree of polymorphism observed among the apple accessions employed in our study compared with that in maize lines, whereby apple has one SNP every 288 bp on average [19], versus every 60 bp in maize [59]. However, it is possible that we might have underestimated the total contribution of CNVs to the apple genome because of the low-coverage sequencing data, strict quality control, and our analysis approach, which focused on detecting only CNVRs that exhibited high variation among the samples.

### In the apple genome CNVRs are enriched in resistance genes

In total, 845 gene models overlapped the CNVRs detected. Initial functional enrichment analysis suggests that these gene models are involved in ion transport, signal transduction, and the defense response. This result is concordant with other studies in a wide range of species, such as humans [4], barley [3], and soybean [10], which have noted that immunity-related genes are frequently found within CNV regions. Fast-mutating CNVs have been recognized as an evolutionary driving force for organisms to adapt to changes in environmental conditions and the introduction of new pests and diseases [60]. This phenomenon is particularly important for long-lived tree species, which have a considerably lengthier generation time than their pathogens and which cannot migrate large distances to adapt to new conditions. In light of this, we propose that CNV is an extremely important adaptive genetic mechanism in tree species, even more so than for other organisms.

### Co-location between CNVRs and trait loci in apple

Numerous studies have focused on identifying genomic regions associated with horticultural traits in apple, using genetic linkage mapping and QTL analysis [see Troggio *et al.* for a review [61]. Three regions of the apple genome that contain putative CNVRs are of particular interest, as they co-locate with loci controlling traits related to biotic and abiotic responses in apple. Firstly, a cluster of 19 CNVRs between 3,507,000 bp and 4,107,000 bp on chromosome 2 contains multiple R genes models (Figures 6 & 7) in a region where major loci conferring resistance to apple scab have been previously mapped. The *Rvi4/Vh4* and *Rvi15/Vr2* loci were originally mapped to apple linkage group 2 [62, 63] and the recent development of genetic markers closely linked to these loci enabled to estimate their physical location on the reference apple genome of ‘Golden Delicious’ [64], close to the detected cluster of 19 CNVRs. Secondly, a CNVR located at positions 3,506,106 and 3,509,841 bp on chromosome 11 co-locates with *Pl2*, a major gene conferring resistance to powdery mildew. *Pl2* was originally mapped using expressed sequence tags-based markers [65] and the newly developed markers for *Pl2* [64] enabled placement of this locus on the apple genome sequence in the same position as the CNVR. Finally, three CNVRs located between positions 1,404,000 and 1,671,000 bp on chromosome 9 co-locate with a QTL for budbreak date [49]. Future work needs to include validation of these CNVRs, to demonstrate linkages between the CNVRs on chromosomes 2, 11, and 9 for scab resistance, powdery mildew resistance and budbreak date, respectively. This validation can be carried out by re-sequencing, with higher depth of coverage, cultivars carrying the scab resistance, powdery mildew, and early budbreak date alleles, and comparing the resulting integer copy-number genotypes to those of cultivars not carrying the resistance. Additionally, using the higher coverage data, the exact breakpoint of an individual CNV can be determined, and in combination with the integer copy-number genotypes, will enable the design of genetic markers that will accurately quantify the copy number at these loci. We hypothesize that some of these CNVs may be causative for these loci and that these functional CNV markers will be more powerful than nearby SNP markers for application in marker-assisted breeding.

## Conclusion

We identified 876 CNVRs with an average size of 16.4 kb, comprising 3.5% of the apple genome. These results represent the first catalogue and investigation of a previously unexplored form of genetic variation in a tree species. The CNVRs identified in this study are enriched in resistance (R) gene models and overlap with major gene loci of agricultural significance. Further investigation of apple CNV using higher coverage NGS data will enable integer-level copy-number assignment and break-point identification. This will facilitate the discovery of the causative CNV and improve the design of molecular markers that segregate with the trait. Ultimately, we believe that the focused investigation of CNV in the apple genome will lead to the genetic improvement of apple cultivars and a deeper understanding of the role CNV plays within the apple genome and other long-lived tree species.

## Acknowledgements

Computational resources from the New Zealand eScience Infrastructure (NeSI) were used for some of the analyses presented here (www.nesi.org.nz). We thank Hoang tan Nguyen (CNVrd2 developer), Tanya Flynn, Anna Gosling, and Murray Cadzow for their useful discussions throughout the course of this project, and Sue Gardiner and Susan Thompson from PFR for useful comments on the manuscript. Funding for this research was obtained from University of Otago, Plant & Food Research’s Core funding and the USDA RosBREED project.

## Additional files

**Additional file 1: Sequencing and quality control result for the 30 apple varieties.**

**Additional file 2: List of copy-number variable regions (CNVRs) identified in the apple genome.**

**Additional file 3: Per chromosome summary of copy-number variable regions (CNVRs) in the apple genome.**

**Additional file 4: List of all gene models found within copy-number variable regions (CNVRs) in the apple genome.**

**Additional file 5: List of resistance (R) gene models located within copy-number variable regions (CNVRs) in the apple genome.**

